# Late cortical potentials are not a reliable marker of somatosensory awareness

**DOI:** 10.1101/2020.10.01.322651

**Authors:** Pia Schröder, Till Nierhaus, Felix Blankenburg

## Abstract

Two types of scalp-recorded event-related potentials have been proposed as neural correlates of perceptual awareness in humans: an early, modality-specific negativity and a late, modality-independent positivity. However, whether these potentials genuinely reflect perception or result from task demands remains controversial. To address this question, we compared results from a classical somatosensory detection task (direct report task) to a somatosensory-visual matching task, in which overt reports were decorrelated from target detection, equated the behavioural relevance of detected and undetected stimuli, and mitigated the influence of attentional processes. By means of Bayesian model selection, we show that the early N140 component was the first to reflect target detection in both tasks, whereas the late P300 component was task dependent, with strong detection effects in the direct report task that were absent in the matching task. We conclude that the P300 is not a genuine correlate of somatosensory awareness but reflects postperceptual processing.

## Introduction

Whether the neural processes giving rise to conscious perception originate in sensory cortices or higher order regions is a long-standing debate. One line of investigation studies event-related potentials (ERPs) recorded from the human scalp. By identifying the latencies at which awareness-related ERPs occur, researchers hope to elucidate the temporal dynamics of conscious perception and infer underlying mechanisms.

In the somatosensory modality, consciously perceived stimuli elicit increased ERP amplitudes starting at ∼90 ms post stimulus (Auksztulewicz, Spitzer, & Blankenburg, 2012; Schubert, Blankenburg, Lemm, Villringer, & Curio, 2006). Most notably, the somatosensory N140 over contralateral somatosensory electrodes and the P300 over centroparietal electrodes have repeatedly been reported to reflect detection of tactile stimuli (Al et al., 2020; Auksztulewicz & Blankenburg, 2013; Auksztulewicz et al., 2012), whereas very early components, such as the P50, seem to reflect physical stimulus properties (Forschack, Nierhaus, Müller, & Villringer, 2020). These findings are consistent with studies in the visual and auditory modalities that have similarly reported awareness-related early negativities at ∼200 ms over modality specific electrodes and late positivities at ∼350 ms over centroparietal electrodes (Eklund & Wiens, 2019; Förster, Koivisto, & Revonsuo, 2020).

Although these potentials are routinely found to correlate with consciously perceived stimuli across various tasks, recent years have seen a rigorous debate centring on the question whether they truly reflect awareness or are instead associated with precursors or consequences of conscious perception (see Förster et al. (2020) for a recent review in the visual modality). Most notably, in a series of studies, Pitts and colleagues have demonstrated that the visual P300 only indexes conscious perception when stimuli are reported, i.e. when they are behaviourally relevant to the task (Cohen, Ortego, Kyroudis, & Pitts, 2020; M. A. Pitts, Metzler, & Hillyard, 2014; Schlossmacher, Dellert, Pitts, Bruchmann, & Straube, 2020). Others have found that the P300 is sensitive to manipulations of attention (Koivisto, Revonsuo, & Lehtonen, 2005), expectations (Melloni, Schwiedrzik, Müller, Rodriguez, & Singer, 2011), and the timing of reports (Ye & Lyu, 2019), altogether suggesting that it might be associated with post-perceptual processing. However, critics of this conclusion have pointed out that failure to detect a P300 for conscious but task-irrelevant stimuli may be due to increased temporal variability of the elicited components (Boncompte & Cosmelli, 2018) or even a failure to correctly identify conscious stimuli considering the difficulty to categorise trials in no-report paradigms (Mashour, Roelfsema, Changeux, & Dehaene, 2020). In contrast, given that sustained sensory negativities are known to be modulated by attention (Eimer & Forster, 2003; Mena, Lang, & Gherri, 2020) it has been suggested that awareness-related negativities reflect precursors of conscious perception and should be scrutinised just as rigorously as the P300 (M. A. Pitts et al., 2014; Rutiku, Martin, Bachmann, & Aru, 2015). The issue is further complicated by the fact that evidence for task dependence of the P300 stems almost exclusively from the visual modality, whereas respective studies are scarce in the auditory (Eklund, Gerdfeldter, & Wiens, 2019) and completely lacking in the somatosensory modality. Taken together, the difficulty to delineate correlates of conscious perception from its precursors and consequences as well as a lack of evidence from modalities other than vision have so far prohibited final conclusions regarding the earliest ERP markers of perceptual awareness.

The goal of the current study was to address these challenges and scrutinise the task dependence of early and late ERP correlates of perceptual awareness in the somatosensory modality. In two experiments, we recorded EEG data from healthy participants performing somatosensory detection tasks that differed in their report requirements. In the first experiment, participants performed a direct report task similar to classical detection tasks. In this experiment, we expected the somatosensory N140 and P300 to correlate with target detection, replicating previous results (Al et al., 2020; Auksztulewicz et al., 2012). In the second experiment, rather than directly reporting hits and misses, participants compared their somatosensory percepts to visual matching cues. The resulting match and mismatch reports were decorrelated from target detection and because detected and undetected targets could result in the same overt report, their behavioural relevance and ensuing working memory was equated. Moreover, the multimodal nature of the task required participants to split their attentional resources between somatosensory and visual inputs and quickly combine the extracted information into corresponding reports. As a result, signal differences related to post-stimulus attentional capture were expected to be minimised. To track the transformation from physical to perceptual processing stages, we constructed simple behavioural models capturing various task dimensions and assessed their ability to explain the EEG signal by means of Bayesian model selection (BMS, Stephan, Penny, Daunizeau, Moran, & Friston, 2009). In a previous study, we employed this approach in combination with functional magnetic resonance imaging (fMRI) and found that detection-related signals were confined to secondary somatosensory cortex (SII), whereas frontal regions reflected stimulus uncertainty and reports (Schröder, Schmidt, & Blankenburg, 2019). Based on these results and previous insights from the visual modality, we hypothesised that (i) very early potentials (P50) reflect physical stimulus properties, (ii) the N140 correlates with target detection independent of task requirements, and (iii) the P300 reflects post-perceptual processes and only differentiates between hits and misses when reports correlate with detection (direct report task) but not when reports, behavioural relevance, and working memory are controlled for (matching task).

## Results

### Experimental paradigm

In both experiments, participants performed two-alternative forced choice somatosensory detection tasks on electrical median-nerve stimuli while their EEG was recorded. The stimulation protocols and visual displays were identical and only the task instructions differed to produce hit/miss reports in the direct report task and match/mismatch reports in the matching task (see **Figure 1A** and Methods for details). On every trial, participants received a target stimulus at one of ten intensity levels. The intensity levels were individually calibrated to sample the full dynamic range of each participant’s psychometric function from 0-100% detectability, such that physical stimulus properties, detection probability, and perceptual uncertainty associated with target detection varied from trial to trial. Simultaneously, a central grey fixation disk changed its brightness to either white or dark grey. In the direct report task, participants were informed that the change to white or dark grey indicated the timing of a potential target stimulus, with no relevance of the direction of change. In the matching task, this change served as the matching cue and participants were instructed to compare their somatosensory percepts (detected vs. undetected) to the matching cue that signalled stimulus presence (white) or absence (dark grey). After a brief delay, participants reported their decision (hit/miss in the direct report task, match/mismatch in the matching task) by making a saccadic eye movement to one of two peripherally presented, colour-coded response cues. Participants could not predict which response cue would be presented on which side so that they could not prepare a motor response early in the trial in either task. However, due to the different report requirements, the two tasks differed in their levels of experimental control. While target detection was expected to correlate with reports, behavioural relevance, working memory, and attentional capture in the direct report task, these variables were controlled for in the matching task (or at least mitigated in the case of attentional capture). Accordingly, the contribution of post-perceptual processing to the hit vs. miss contrast was expected to be considerably attenuated.

**Figure 1.**
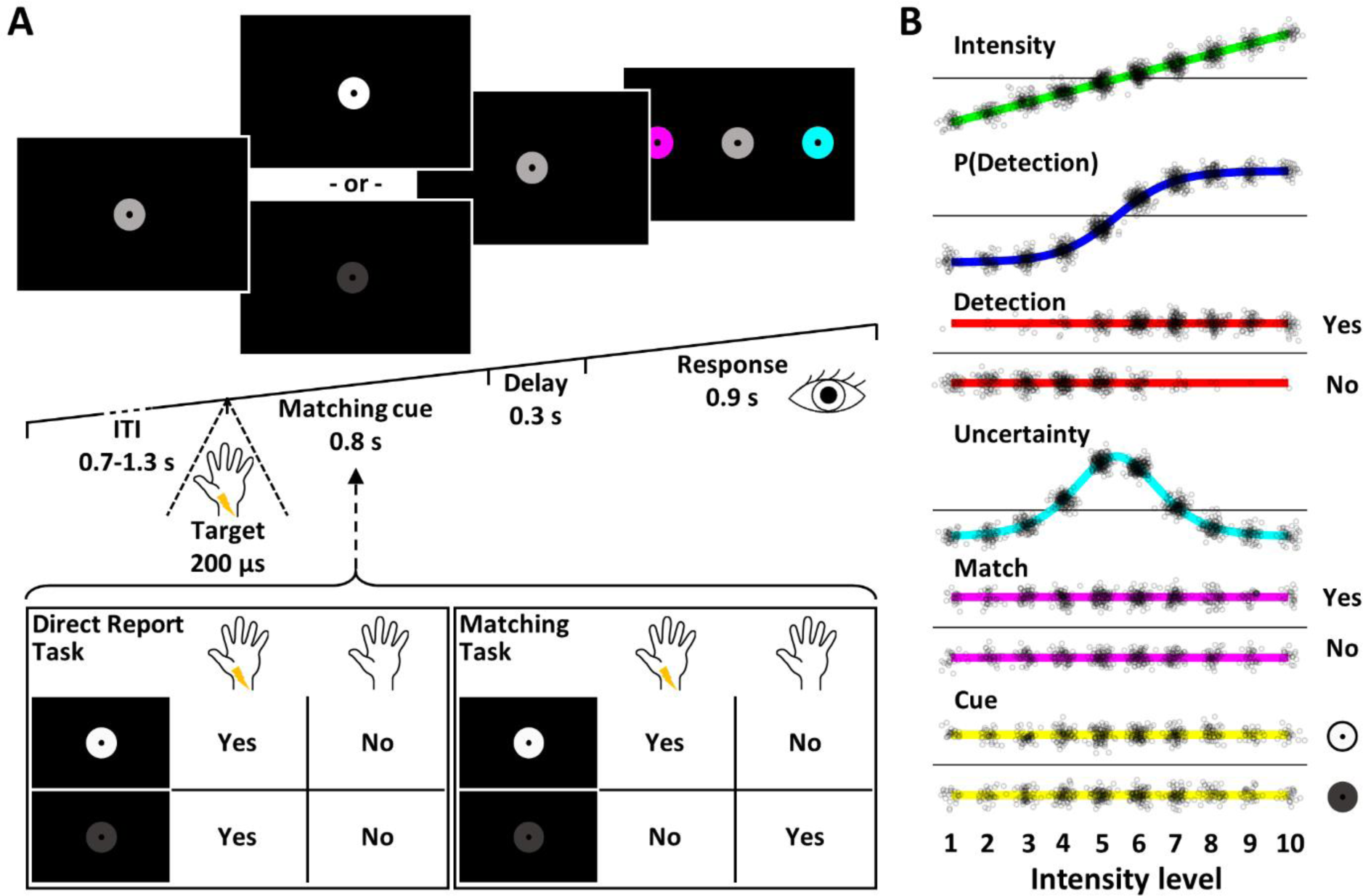
Experimental Design. **(A)** Trial design. Following a variable intertrial interval, participants received an electrical target pulse at one of ten intensity levels, which they either detected or missed. At the same time, the fixation disk changed its brightness to serve as a visual matching cue that signalled stimulus presence (white) or stimulus absence (dark grey). In the direct report task, participants ignored the matching cue and merely decided whether they had detected a target pulse or not (left inset). In the matching task, participants compared their somatosensory percept to the matching cue and decided whether the two modalities matched or not (right inset). After a brief delay, they reported their decision by making a saccade to one of two peripherally presented, colour-coded response cues. The selected cue briefly increased in size, signalling that the response was logged and the next trial began. **(B)** Experimental Regressors. EEG responses were modelled with seven different GLMs that were compared using BMS. Each experimental GLM contained an intercept regressor and one of six experimental regressors modelling stimulus intensity, detection probability, target detection (hit vs. miss), expected uncertainty, matching reports (match vs. mismatch), and matching cues (white vs. dark grey). An additional null model contained the intercept regressor only. Small black circles denote individual trials of one participant. Note the intensity-biased distribution of trials in the detection regressor, leading to correlations between models, which prohibit classical GLM analysis.

### Behaviour

Participants detected 55.84 ± 8.99% of trials in the direct report task, and 53.77 ± 9.49% in the matching task. Target detection was most variable on trials with intermediate stimulus intensity levels resulting in characteristic sigmoidal psychometric curves (**Figure 2**). Reaction times were slightly shorter for hits than misses (direct report task: hits: 313.62 ± 34.17 ms, misses: 333.89 ± 42.28 ms, Bayes factor in favour of a difference (BF10) = 568.68; matching task: hits: 308.97 ± 32.25 ms, misses: 314.05 ± 36.52 ms, BF10 = 9.27). For the matching task, we used Bayesian tests of association to test if the task successfully dissociated target detection from overt reports and found that target detection and matching reports were indeed independent (Bayes factor in favour of the null hypothesis of independence (BF01) > 5 for all participants).

**Figure 2.**
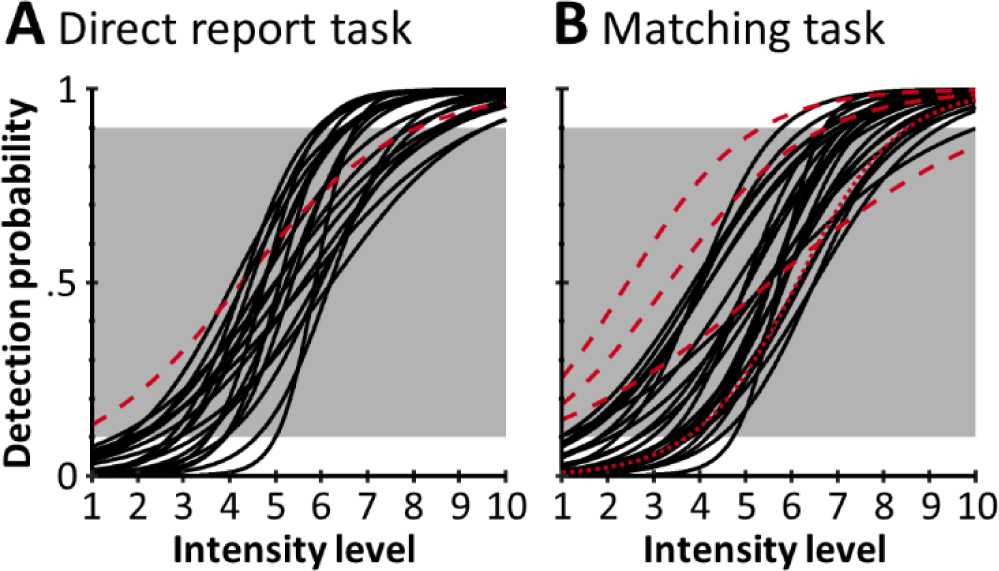
Psychometric functions in the direct report task (**A**) and the matching task (**B**). Black lines correspond to the individual block-averaged psychometric functions of participants included in the final samples (direct report task: n = 22; matching task: n = 24). Red dashed lines correspond to participants whose detection probabilities at minimum and maximum intensity levels fell outside the required margin of <10% and >90% (white background) and were thus excluded from the analyses. The red dotted line corresponds to one participant that was excluded due to poor EEG data quality.

### EEG

Neuronal processing of the somatosensory stimulus was expected to undergo a transformation from physical to perceptual stages and on to the final reports. To track this transformation in the EEG signal and identify the latency at which awareness-related potentials occur, we constructed five general linear models (GLMs). Each contained an intercept regressor and a trial-wise experimental regressor capturing a variable of interest: 1. the linear stimulus intensity as a model of physical stimulus properties, 2. the individual psychometric functions modelling detection probability, 3. binary target detection modelling a non-linear response expected to index awareness, 4. the slope of individual psychometric functions modelling the expected uncertainty associated with target detection, and 5. match and mismatch reports (**Figure 1B**). Note that even though participants did not engage with the matching cue in the direct report task and did not form match/mismatch reports, we still included the match model (as defined by the alignment of participants’ hit/miss reports with the matching cues presented on every trial) in the analysis of both experiments to ensure that any differences in results would be attributable to differences in the data and not to differences in the model spaces. In addition to the five models of interest, we included two control models to validate the approach. A visual control model was defined by the matching cues (white vs. dark grey) to demonstrate specificity of the results to electrodes over somatosensory and visual cortices, respectively, and a null model contained only the intercept regressor to ensure that model selection would correctly dismiss experimental models when no effects were expected (i.e. no effects in the baseline period).

We fitted the GLMs to participant’s trial-wise EEG data using the Bayesian estimation scheme as implemented in SPM12 to obtain estimates of log model evidence (LME) per model, participant, electrode, and time point. To identify which model best explained the EEG data at every time point, the estimated LMEs were then used to perform time-resolved BMS. BMS computes exceedance probabilities (EPs) for all models, quantifying the probability that a particular model explains the data better than any of the other models. In addition to allowing for model comparison of non-nested models, this approach offers the possibility to combine similar models into model families and assess them on the family level (Penny et al., 2010), which facilitates fair comparison even when some of the included models are correlated and thus, share probability mass. In our case, because target detection becomes more likely with increasing stimulus intensity, the intensity, detection probability, and detection models were expected to correlate positively. Therefore, we combined these models into a model family (which we termed +family due to their positive correlations) and performed BMS on the family level, comparing the +family with the uncertainty, match, cue, and null models. The individual +family models were then further assessed only at those time points that were well explained by the +family.

To identify time points showing significant effects, we imposed two criteria: 1) the winning model family had to score an EP ≥ .99 and 2) across participants the beta estimates of the winning models’ experimental regressors had to significantly deviate from zero to ensure a systematic effect across participants. Significant time points were defined as those exceeding both the EP threshold and a beta evidence threshold of BF10_beta_ ≥ 3. The null model was exempt of this rule as it did not have an experimental regressor and thus was only required to exceed the EP threshold.

#### Task dependence of early and late somatosensory ERPs

To test our hypotheses regarding the task dependence of the P50, N140, and P300, we inspected the BMS results in three electrodes of interest: CP4, C6, and CPz **(Figure 3)**. The grand-averaged EEG signals plotted for each intensity level showed the largest deflections for stimuli of high intensity levels and the smallest deflections for stimuli of low intensity levels, suggesting correlation with the +family. This observation was confirmed by BMS: In both tasks, the P50 in contralateral electrode CP4 was best explained by the intensity model, indicating processing of physical stimulus properties at this latency. The P50 was followed by a centroparietal P100, which was similarly modulated by stimulus intensity in both tasks but showed a transition to detection probability at ∼120 ms in the matching task. The N140 in electrode C6 was modulated by target detection in both tasks, but in the matching task this effect occurred only after the component had peaked, at ∼150 ms, and was preceded by an effect of stimulus intensity. Finally, the P300 in electrode CPz showed clear task dependence. In the direct report task, the early phase of the P300, starting at ∼200 ms, was best explained by the intensity model, whereas its later phase, starting at ∼350 ms, showed a sustained effect of target detection. This transition was apparent in most centroparietal electrodes **(Supplementary Figure 2)**, suggesting a broad distribution of the effects. In contrast, in the matching task, from ∼250 ms onwards the P300 in electrode CPz was best explained by the detection probability model, with no effects of target detection. Here, we found different response profiles in different centroparietal electrodes: contralateral electrodes showed primarily intensity effects, midline electrodes tended to show detection probability effects, and two ipsilateral electrodes (C3 and CP3) showed brief detection effects at ∼320 ms (**Supplementary Figure 3)**.

**Figure 3.**
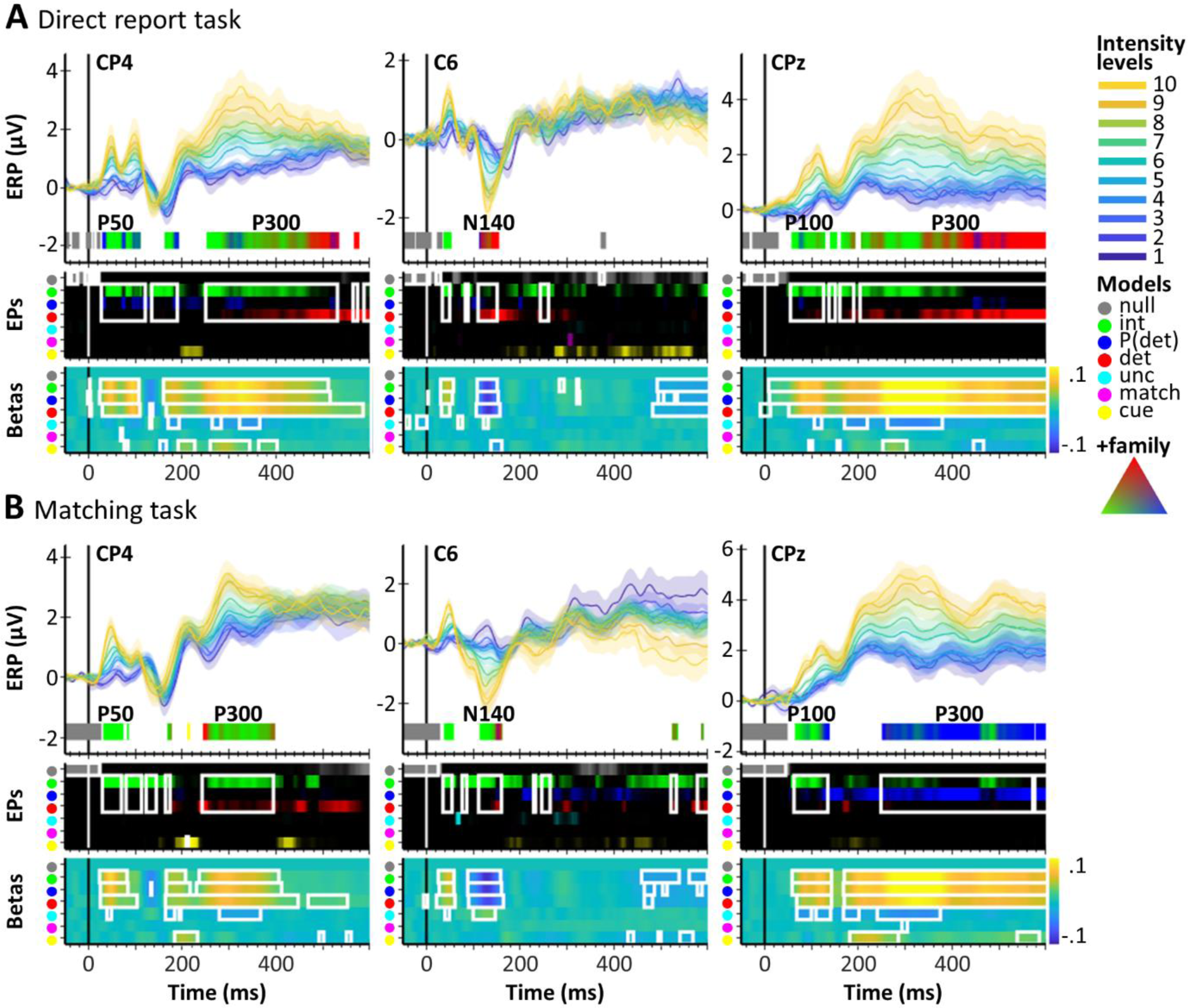
ERPs and BMS results for three electrodes of interest (CP4, C6, CPz) in the direct report task (**A**) and in the matching task (**B**). For each electrode, the top panel shows stimulus-locked, grand-averaged ERPs (mean ± standard error) for each intensity level (1-10). Below, BMS results are plotted for significant time points (EP ≥ .99 and BF10_beta_ ≥ 3). The colour bands indicate the winning model families. For time points best modelled by the +family (intensity, P(detection), detection), the colour corresponds to an RGB value that is composed of the EPs of the three +family models (compare the RGB triangle: corners correspond to EP = 1, signifying a clear winner of the model comparison within the +family, whereas intermixed colours signify similar EPs for the respective models). The middle panel shows unthresholded EP time courses for each model. The bottom panel shows time courses of group-averaged beta estimates of each model’s experimental regressor (warm colours denote positive beta estimates, cold colours denote negative beta estimates). White rectangles mark data segments that exceed the respective thresholds. The results suggest that the P50 was modulated by stimulus intensity in both tasks. The N140 showed and effect of target detection in the direct report task and a transition from stimulus intensity to target detection in the matching task. The P300 was strongly task dependent, showing a transition from stimulus intensity to target detection in the direct report task, and an effect of detection probability in the matching task.

To further inspect the different detection effects in the two tasks, we plotted hit and miss ERPs that were matched for intensity levels (see **Supplementary Figure 1** for details). The resulting grand-averaged ERPs showed no difference between hits and misses in the P50 component, further confirming that early EEG signals reflect processing of physical stimulus properties (**Figure 4**). The N140 on the other hand showed a clear amplitude enhancement for hits compared to misses and this effect was apparent in both experiments, even though in the matching task, the detection model only dominated the model comparison in a later time window. In contrast, the P300 showed a strong amplitude enhancement for hits compared to misses in the direct report task but not in the matching task. In both tasks, the signal first increased at ∼200 ms for both hits and misses. However, in the direct report task, this signal increase was much stronger for hits than misses, whereas in the matching task, hit and miss amplitudes were comparable and this was true even for electrodes showing brief detection effects (compare **Supplementary Figures 4 and 5**). Note that although this way of looking at the results did not involve any further statistical testing, it helps to better understand the evolution of detection effects in the data. E.g. looking at the intensity-matched hit and miss ERPs in electrode CPz, the late detection effect in the direct report task appears to emerge as early as 200 ms (compare **Figure 4A**). However, the BMS results indicate that the detection effect only started at ∼350 ms whereas earlier time points where explained by stimulus intensity. This suggests that although target detection seems to have had some effect on the signal already at 200 ms, the intensity model still did a better job at explaining the data at this point and only later, at ∼350 ms, did the detection model become the most dominant. Similar conclusions may be drawn for the early phase of the N140.

**Figure 4.**
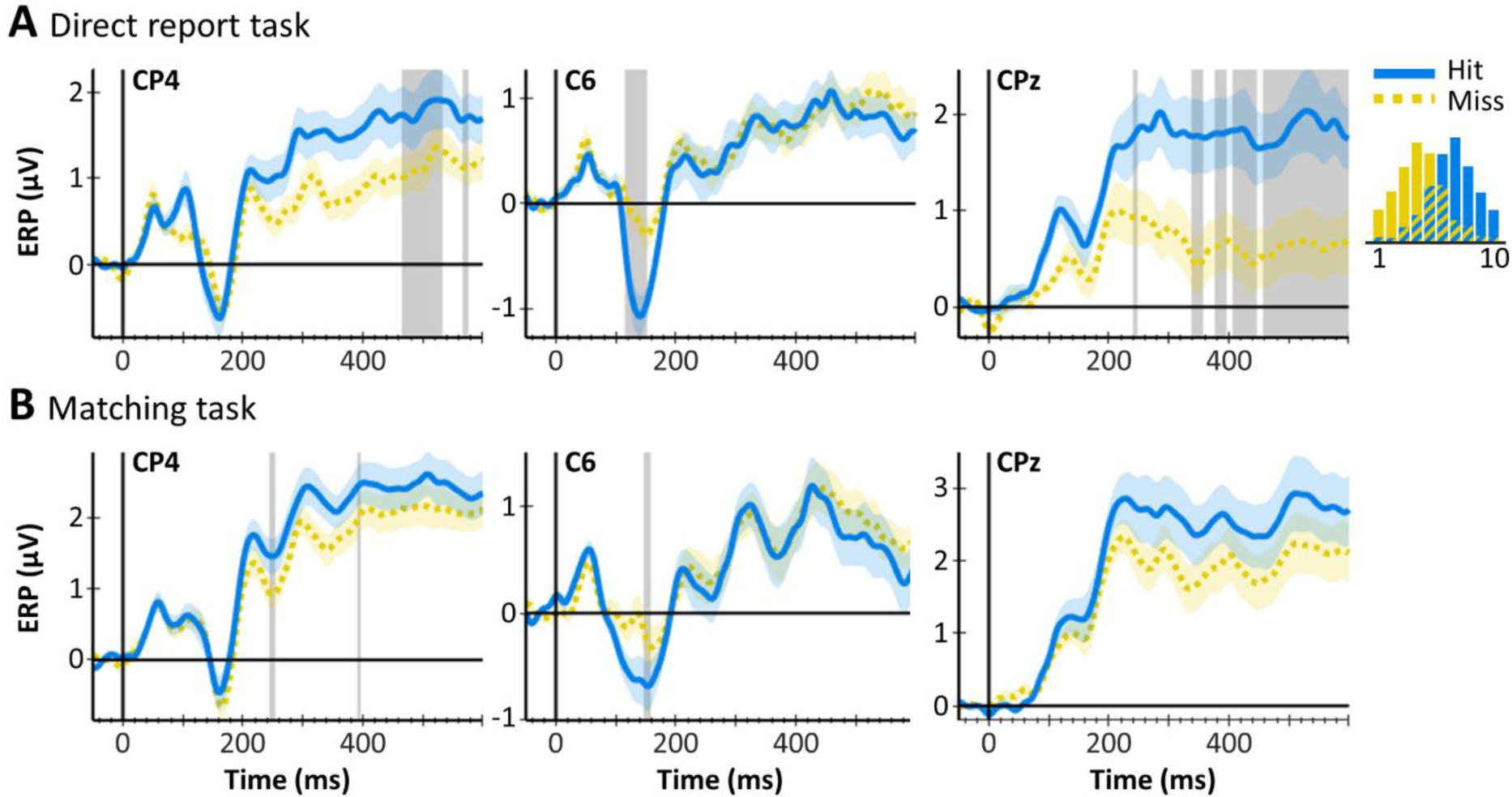
Intensity-matched hit and miss ERPs (mean ± standard error) in the direct report task (**A**) and in the matching task (**B**). Grey shaded areas mark time points that were best explained by the detection model. The histogram on the right shows trial distributions for hits (yellow) and misses (blue) across intensity levels and the overlap indicates the intensity-matched subsample (for details see **Supplementary Figure 1**). The P50 was not modulated by target detection in either task, whereas the N140 exhibits larger amplitudes for hits compared to misses in both tasks. The P300 shows a large difference between hits and misses in the direct report task but not in the matching task.

#### Effects across time and space

To obtain a more global impression of the spatiotemporal evolution of model probabilities, we determined the overall model performances across time as defined by the proportion of electrodes showing significant effects for each model (**Figure 5**). As expected, activity in the baseline period was best explained by the null model, validating the specificity of our analysis approach. The intensity model showed similar effects in both tasks: the first clear evidence of somatosensory stimulus processing emerged at ∼50 ms, consistent with the P50 response. The BMS topographies at this time point showed a wide-spread intensity effect in contralateral somatosensory and frontal electrodes, which rotated slightly throughout the following ∼100 ms to include more centroparietal electrodes. Then, starting at ∼250 ms, a second strong intensity effect occurred, encompassing primarily centroparietal electrodes. The detection probability model showed several smaller peaks in the direct report task with similar topographies as the early intensity effect, but across time and electrodes this model did not explain the data well. In the matching task on the other hand, we found a detection probability effect at ∼120 ms and an additional late effect starting at ∼250 ms that paralleled the late intensity effect. Both of these effects showed a centroparietal topography, but the intensity effect occurred primarily in contralateral electrodes, whereas the detection probability effect was mostly found in slightly more posterior midline and ipsilateral electrodes. The detection model showed the most striking difference between the two tasks. In the direct report task, the earliest detection effect occurred at ∼90 ms in ipsilateral temporal electrodes. A more widespread detection effect in contralateral central and frontal electrodes followed at ∼120 ms (including the N140) and yet another effect occurred at ∼250 ms in ipsilateral central electrodes. Finally, from ∼350 ms onwards, a large centroparietal electrode cluster reflected target detection throughout the rest of the time window. These detection effects were greatly reduced in the matching task. Here, the earliest effect was observed at the N140 latency and was confined to only two electrodes (C6 and FC6) and a brief period of time. Most strikingly, the wide-spread centroparietal detection effect at ∼350 ms observed in the direct report task was virtually absent in the matching task. Instead, centroparietal signals in the P300 time range were best explained by the intensity and detection probability models, with only very brief effects of target detection in isolated electrodes (**Supplementary Figure 3**). Surprisingly, neither the uncertainty nor the match model scored high EPs in either task and this lack of effects was unaltered by inspecting later time points or relaxing the significance criteria. However, we found effects of the matching cue in both tasks, starting at ∼90 ms in occipital electrodes. In the matching task, this effect lasted slightly longer (until ∼300 ms) and encompassed frontal and parietal electrodes, indicating further processing. This difference was unsurprising given that processing of the matching cue was vital to the matching task but not to the direct report task.

**Figure 5.**
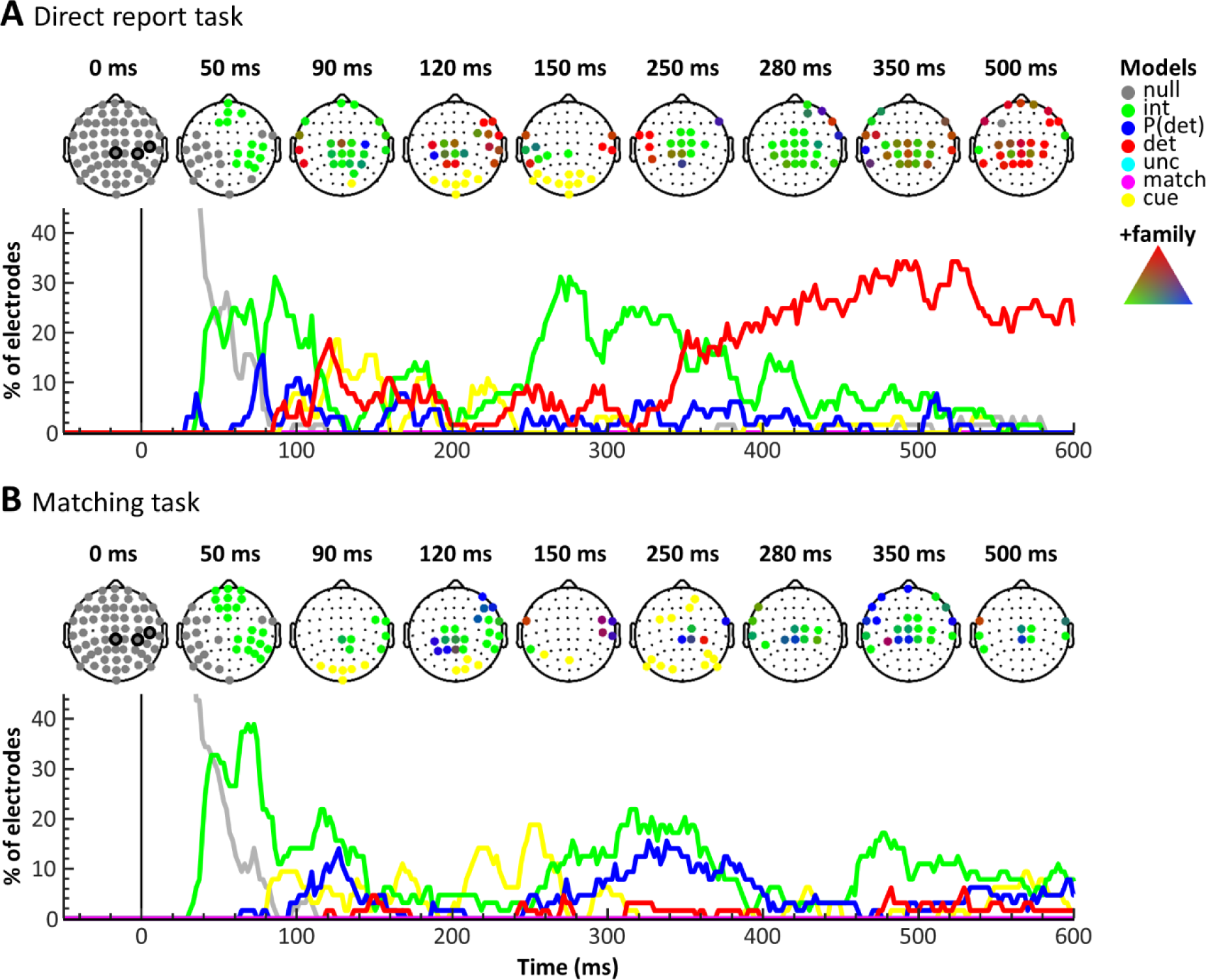
BMS results across electrodes in the direct report task **(A)** and the matching task **(B)**. For each task, scalp topographies for time points of interest (top) and model time courses across electrodes (bottom) are displayed. The scalp topographies indicate winning models in significant electrodes using colours as in Figure 3. The circled electrodes at 0 ms mark electrodes CP4, C6, and CPz. Model time courses are plotted as the proportion of electrodes showing significant effects over time. The results suggest a striking reduction of target detection effects (red) in the matching task compared to the direct report task, especially in late time windows >350 ms.

## Discussion

In this study, we scrutinised the relevance of early and late ERP components as markers of somatosensory awareness. Both in a classical detection task with direct reports and in a revised version that used a matching procedure to control for report requirements, the somatosensory P50 component was modulated by physical stimulus intensity and showed no sensitivity to target detection, confirming hypothesis (i). In contrast, the N140 was modulated by target detection in both tasks, although in the matching task, this modulation occurred later and was preceded by an effect of stimulus intensity, such that hypothesis (ii) was only partly confirmed. The commonly reported late detection effect in the P300 time window was strongly task dependent. While in the direct report task, we observed a widespread signal increase for hits compared to misses that was best explained by target detection from ∼350 ms onwards, this effect was largely absent in the matching task, when report requirements were controlled for. These results confirmed hypothesis (iii) and suggest that the P300 is not a reliable marker of perceptual awareness but reflects postperceptual processing.

The intensity effect observed in the P50 component replicates a recent ERP study that has similarly shown scaling of the P50 amplitude with stimulus strength (Forschack et al., 2020). More generally, our finding is in line with research showing that early stimulus processing in SI does not differentiate between detected and undetected stimuli but reflects physical stimulus properties in both humans (Schröder et al., 2019; Schubert et al., 2006) and macaques (de Lafuente & Romo, 2005, 2006). We thus add to the growing consensus that very early sensory potentials index preconscious processing (Förster et al., 2020) and support the notion that initial feedforward processing in early sensory regions is not sufficient for conscious perception.

The N140 component has repeatedly been reported as a correlate of tactile awareness (Al et al., 2020; Auksztulewicz & Blankenburg, 2013; Auksztulewicz et al., 2012; Forschack et al., 2020; Schubert et al., 2006; Zhang & Ding, 2009). Likewise, in our study, the earliest effect of target detection common to both tasks was observed at the N140 latency. Similar to the visual awareness negativity (VAN, Koivisto & Grassini, 2016) and auditory awareness negativity (AAN, Giani, Belardinelli, Ortiz, Kleiner, & Noppeney, 2015), the N140 has been suggested to result from recurrent interactions between sensory cortices (Auksztulewicz & Blankenburg, 2013; Auksztulewicz et al., 2012). Accordingly, these early negativities have been taken as evidence for the Recurrent Processing Theory (RPT) of conscious perception, which assumes that perceptual awareness emerges as soon as initial feedforward activity in sensory cortices is consolidated by re-entrant feedback (Lamme, 2006). In somatosensation, a number of studies have demonstrated detection-related feedback signals from SII to SI in humans (Auksztulewicz & Blankenburg, 2013; Auksztulewicz et al., 2012; Jones, Pritchett, Stufflebeam, Hämäläinen, & Moore, 2007), mice (Kwon, Yang, Minamisawa, & O’Connor, 2016; Yang, Kwon, Severson, & O’Connor, 2016) and macaques (Cauller & Kulics, 1991), supporting the role of local recurrent processing for somatosensory awareness. Interestingly, using dynamic causal modelling (Friston, Harrison, & Penny, 2003), Auksztulewicz and colleagues (2012) found that a recurrent SI-SII model first outperformed a purely feedforward model at 160 ms. This transition aligns well with the transition from stimulus intensity to target detection observed in the N140 component in our study. Taken together, these results may suggest that physical stimulus properties are signalled in a feedforward manner whereas perceptual properties are conveyed in feedback pathways, a perspective consistent with RPT.

Notably, while we did find larger N140 amplitudes for hits compared to misses in both tasks, the BMS results revealed a delayed onset and reduced spatial extent of the detection effect in the matching task. This difference suggests that even though the N140 was not abolished by controlling for report requirements, the different task demands did have a considerable influence on its timing and distribution. In fact, the N140 is well known to be modulated by endogenous (Eimer & Forster, 2003) and exogenous attention (Mena et al., 2020). Both of our tasks varied intensity levels, which reduces the impact of prestimulus attentional focus on perceptual outcome and thus, we expected the effects of endogenous attention to be negligible compared to near-threshold detection tasks. However, in contrast to the direct report task, the matching task required participants to split their attentional resources between the somatosensory and visual modalities and immediately transform the extracted information into appropriate reports. Accordingly, post-stimulus attentional engagement was expected to be alleviated in the matching task, which may be the reason for the reduced N140 detection effect. Awareness and attention are closely related concepts that are difficult to separate both conceptually and experimentally (Koch & Tsuchiya, 2012). Nonetheless, our current results suggest that at least part of the detection effects routinely reported at the N140 latency may be due to uncontrolled attentional processes and more direct manipulations of attentional allocation should be employed to further elucidate this issue.

The P300 has repeatedly been demonstrated to depend on report requirements in visual tasks (Cohen et al., 2020; M. Pitts, Padwal, Fennelly, Martinez, & Hillyard, 2014; Schlossmacher et al., 2020) and we show that the same is true in the somatosensory domain. Crucially, in contrast to previous studies our approach did not rely on making stimuli irrelevant or removing reports and thus, our results cannot be explained in terms of increased signal variability or falsely labelled trials. Instead, we show that by altering report requirements, a P300 was elicited for both hit and miss trials, highlighting that it is not contingent on perceptual awareness and thus, unlikely to constitute a genuine correlate of conscious perception. In opposition to early negativities, the P300 has been taken as evidence in support of the Global Neuronal Workspace Theory (GNWT). GNWT assumes that any conscious percept presupposes a non-linear “ignition” of global workspace neurons that are distributed across a network of brain regions, primarily in frontal and parietal cortices (Baars, 2002; Dehaene & Naccache, 2001). Given that late activity is considered a hallmark of ignition (Dehaene & Changeux, 2011), evidence for task dependence of the P300 constitutes a considerable challenge to the GNWT. It has recently been argued that disqualifying the P300 as a correlate of awareness does not disqualify cognitive or higher order theories of awareness (Cohen et al., 2020) and we agree that demonstrating the task dependence of the P300 alone does not prove the absence of ignition. However, numerous studies using different tasks and imaging methods have demonstrated that activity in frontal and parietal regions is equally task dependent (Farooqui & Manly, 2018; Frässle, Sommer, Jansen, Naber, & Einhäuser, 2014; Schröder et al., 2019), and the apparent absence of any sign of ignition in these studies has not convincingly been addressed by GNWT’s proponents. Importantly, the explanation that early activity may reflect “information accessibility”, whereas late activity reflects “conscious access” and therefore requires some form of reporting (Mashour et al., 2020) does not hold with regards to our results. In both tasks, participants gave reports on every trial and thus conscious access was ensured. One might argue that the high postperceptual demands imposed by the matching task may have concealed detection-related effects in the P300. However, proponents of the GNWT clearly state that perceptual awareness elicits a non-linear ignition, i.e. a large response that is fundamentally different from unconscious processing (Mashour et al., 2020). We would expect such a response to be detectable in late EEG signals, regardless of the added postperceptual demands related to matching reports. Thus, our results do not provide evidence for global ignition as a correlate of somatosensory awareness.

Given the ubiquity of the P300 in studies on perceptual awareness, it is worth considering which cognitive variables are its most likely generators and should therefore be most rigorously controlled. The large number of studies reporting P300 effects for many different cognitive processes suggests a certain heterogeneity in its functional role but a recent review by Verleger (2020) argues that, among the different potential processes, stimulus-response-link reactivation, memory storage, and closure of cognitive epochs are the most relevant. Both our tasks provided timing cues that indicated the exact moment of a potential stimulus delivery and thus, controlled for closure of cognitive epochs. However, the two tasks clearly differed in their control of stimulus-response-link reactivation and memory storage, which may have caused the different results. In the series of no-report experiments conducted by Pitts and colleagues (Cohen et al., 2020; M. A. Pitts et al., 2014; Schlossmacher et al., 2020), the stimuli of interest were not relevant to the task, such that none of the processes in question were elicited upon stimulus perception and no P300 was elicited. In contrast, Sanchez and colleagues showed late decoding of target detection in a go/no-go paradigm with response reversals (Sanchez, Hartmann, Fuscà, Demarchi, & Weisz, 2020) but their task was not controlled for closure or working memory effects. Using a similar task, Eklund and colleagues (2019) tested the task dependence of the auditory P300 and while they did control for closure by providing timing cues, the memory confound likely remained. Thus, it remains unclear whether the reported P300 effect truly reflects awareness, further disqualifying the go/no-go approach as a proper control for task demands (but see Koivisto, Salminen-Vaparanta, Grassini, & Revonsuo, 2016). It becomes clear that in order to effectively further our understanding of the P300’s relevance for perceptual awareness, future studies should carefully control for concomitant processes that are known to produce signals in the P300 time range, in particular stimulus-response-link reactivation, memory storage, and closure of cognitive epochs, such that observed effects can unequivocally be attributed to stimulus awareness.

Overall, the matching task in our study lead to a drastic reduction of detection effects. While this is in line with our fMRI results, where the detection effect was restricted to small subregions within SII (Schröder et al., 2019), it is somewhat puzzling that the conscious percept of a stimulus, an event that is often described as a non-linear, threshold-crossing event (Dehaene & Changeux, 2011; Del Cul, Dehaene, Reyes, Bravo, & Slachevsky, 2009; Mashour et al., 2020; Sergent & Dehaene, 2004), should result in such restricted brain activation. An interpretation that may account for this apparent lack of signals reflecting perceptual outcomes, considers the subjective dimension of the task. Even though the intensity of the stimuli was not directly relevant to participants’ behaviour, we can presume that their subjective experience of a detected target at intensity level 5 differed considerably from the experience of a target at intensity level 10. Importantly, such differences in subjective experience would be expected to covary with the intensity and detection probability models at least for detected stimuli and indeed, we found widespread effects of intensity and detection probability in the P300 time range. The goal of our task design (decorrelation of target detection and overt reports) prohibited subjective awareness ratings, which could give more direct insights into this matter, but previous research indeed suggests that subjective experience of low-level perceptual features may have a graded rather than dichotomous, all-or-none nature (Overgaard, Rote, Mouridsen, & Ramsøy, 2006; Tagliabue, Mazzi, Bagattini, & Savazzi, 2016). In fact, a previous study in the somatosensory modality has demonstrated that the subjective experience of electrical targets of identical stimulus intensity can vary considerably and that this variation correlates with the amplitudes of the N140 and P300 (Auksztulewicz & Blankenburg, 2013). On the other hand, a study on visual awareness has reported that while the VAN scaled with subjective awareness levels, the P300 showed a categorical response (Sergent, Baillet, & Dehaene, 2005). However, due to the direct response mappings used in this study, it remains unclear whether this result genuinely reflects dichotomous perception or followed from task demands. Still, the graded perception view may potentially rehabilitate the relevance of late EEG signals for subjective experience. It is important to note, however, that in our fMRI study, intensity and detection probability effects were exclusively found in primary and secondary somatosensory regions. We can only speculate, whether the late effects observed in the EEG experiment were also generated in these regions or instead reflect activity that was not amenable to fMRI. The former interpretation is supported by findings demonstrating that SII exhibits tonic activity that lasts >300 ms following median nerve stimulation (Avanzini, Pelliccia, Lo Russo, Orban, & Rizzolatti, 2018) and the fact that the P100 component, which is believed to originate in SII (Hämäläinen, Kekoni, Sams, Reinikainen, & Näätänen, 1990; Yamashiro et al., 2019), shows a centroparietal scalp topography that strongly resembles that of the late intensity and detection probability effects observed in our study. Sanchez and colleagues (2020) have reported that target detection can be decoded from sensory cortical regions as late as 350-500 ms poststimulus, further supporting the notion that late activity in sensory regions may reflect perceptual contents. Taken together, the late effects observed in the matching task may have been driven by signals reflecting graded perceptual experience and potentially originate from somatosensory regions. Note however, that this interpretation is entirely speculative at this point and further research using explicit measures of subjective experience will be necessary to scrutinise this proposal.

Although the P300 in our study was clearly affected by report requirements, it did not correlate with reports in the matching task. In fact, the match and uncertainty models did not explain any segments of the EEG data in either task. Possibly, the anatomical locations of regions previously found to show such effects (on the medial wall inside the longitudinal fissure or folded deeply into the cortex, Schröder et al., 2019) prevented strong influences on the overall EEG signal, leading the respective effects to go unnoticed or, alternatively, signals related to uncertainty and matching reports may not have been time-locked to the stimuli. Whatever the reason, since both perceptual uncertainty and overt reports were accounted for in the matching task, we are confident that the observed detection effects were free of these potential confounds (see Schröder et al. (2019) for a more extensive discussion of this point).

In conclusion, our study adds to the accumulating evidence suggesting that the P300 reflects postperceptual processes and is not a genuine correlate of perceptual awareness. We complement this debate with evidence from the somatosensory modality and address previous concerns regarding the reliance on no-report paradigms. More generally, the limited effects of dichotomous target detection in both our EEG and fMRI studies speak against global, non-linear mechanisms of conscious perception. Thus, focussing on large signal divergences for perceived versus unperceived stimuli may be misleading as they seem to emphasise postperceptual processing rather than perceptual awareness. Instead, exploring alternative approaches that probe graded subjective experience while controlling for report requirements may prove a promising avenue for future research.

## Methods

### Participants

All participants reported to be healthy with no history of neurological or psychiatric disorders, were right-handed, and had normal or corrected-to-normal vision. Twenty-three participants completed Experiment 1 (direct report task). Data of one participant were excluded because they did not show a stable psychometric function (see Behavioural analysis and **Figure 2**), leaving data of 22 participants that entered the analyses (twelve females, ten males, age range: 21-42). Twenty-eight participants completed Experiment 2 (matching task). Data of three participants were excluded due to unstable psychometric functions and data of another participant were excluded due to strong motion-related artefacts in the EEG recordings leading to the exclusion of >50% of trials. Thus, data of 24 participants entered the analyses (19 females, five males, age range: 19-37). All participants gave written informed consent prior to the experiment and received a monetary reimbursement or course credits for their participation. The study was approved by the local ethics committee at the Freie Universität Berlin and complied with the Human Subjects Guidelines of the Declaration of Helsinki.

### Experimental paradigm and procedure

The two experiments employed identical visual and somatosensory stimulation protocols. Target pulses were generated as analogue voltage signals using a waveform generator (DT-9812, Data Translation, Bietigheim-Bissingen, Germany), converted to direct current monophasic square wave pulses of 200 µs duration by a constant current stimulator (DS5, Digitimer, Hertfordshire, UK), and delivered via adhesive electrodes (GVB-geliMED GmbH, Bad Segeberg, Germany) attached to the left wrist to stimulate the median nerve. Each trial began with a variable interstimulus interval (ITI, 0.7-1.3 s), during which participants had to fixate on a central grey fixation disk (**Figure 1A**). Fixation was ensured using an SMI RED-m remote eye tracker (120 Hz, Sensomotoric Instruments GmbH, Teltow, Germany) and had to be maintained for at least 0.3 s for a trial to proceed. The stimulation period began with an electrical pulse administered at one of ten intensity levels, which participants either detected or missed. The intensities for each participant were individually determined prior to the experiment to cover the full dynamic range of each participant’s psychometric function from 0-100% detection probability (1%, 50%, and 99% detection thresholds: direct report task: T01 = 2.38 ± 0.78, T50 = 2.88 ± 0.82, T99 = 3.38 ± 0.91; matching task: T01 = 1.88 ± 0.79, T50 = 2.53 ± 0.75, T99 = 3.18 ± 0.91 [mean ± standard deviation]). Individual psychometric functions were estimated using the threshold estimation procedure described in Schröder et al. (2019), which accommodates between-subject variation in detection thresholds and criteria. Presentation of the target pulse was accompanied by a simultaneous change in the fixation disk’s brightness, which served as a timing cue in the direct report task and as the visual matching cue in the matching task. The disk turned either white, signalling stimulus presence, or dark grey, signalling stimulus absence and was presented for 0.8 s. In the direct report task, participants were instructed to ignore the direction of the brightness change and merely decide whether at the time of change they had detected an electrical target or not (**Figure 1A** left inset), such that overt reports directly reflected participants’ perception. In the matching task, participants compared their somatosensory percepts (electrical pulse detected or not detected) against the visual matching cue (signalling stimulus presence or absence) and decided whether the two modalities produced a match (e.g. electrical pulse detected and white matching cue presented) or a mismatch (e.g. electrical pulse detected and dark grey matching cue presented, **Figure 1A** right inset). Note that electrical pulses were presented on every trial and only their intensity levels varied, rendering them sub- or supraliminal. Target detection for the highest and lowest intensities was expected to be relatively stable (∼0% detection at intensity level 1 and ∼100% detection at intensity level 10), whereas detection at intermediate intensity levels was expected to fluctuate from trial to trial. In contrast, the two brightness levels of the matching cue were clearly discernible. This ensured that even in the matching task, electrical target detection could be directly inferred from the combination of the matching cue presented on a trial and the participant’s match or mismatch report. After a brief delay of 0.3 s, in which the fixation disk returned to its original brightness, two colour-coded response cues were presented to the left and right of the fixation disk. Participants reported their hit/miss or match/mismatch decisions by making a saccade to the corresponding response cue (the colour-codes were counterbalanced across participants). Their gaze was evaluated online, and a response was registered as soon as the gaze remained within the response area for 0.2 s, which was signalled to the participant by a brief increase in the selected response cue’s size. The next ITI began as soon as a response was logged or once the allowed response time of 0.9 s was reached. In this case, the fixation disk briefly turned red signalling a missed trial. The presented matching cues and the specific sides on which the colour-coded response cues were presented were counterbalanced across intensity levels, and trials were presented in random order. This procedure resulted in a decorrelation of target detection, matching cues, and motor responses in both tasks, and of target detection and reports in the matching task. Moreover, the selected intensities ensured an overall detection rate of ∼50%, such that detected stimuli were not expected to produce oddball effects. All participants completed 6 blocks of 200 trials (∼10 min per block), resulting in 1200 trials per participant. The number of trials per intensity level followed a normal distribution, such that the number of trials at intensity levels near the 50% detection threshold was maximised (with 192 trials each for the threshold intensity levels 5 and 6 and 48 trials each for the lowest and highest intensity levels 1 and 10). Prior to the matching task experiment, participants completed a 30 min training to ensure full understanding of the task. Stimulus presentation and response collection were implemented in Matlab (The MathWorks, Inc., Natick, MA, RRID:SCR_001622) using the Psychophysics toolbox (Brainard, 1997, RRID:SCR_002881) and SMI’s iView X SDK (Sensomotoric Instruments GmbH, Teltow, Germany).

### EEG recording and preprocessing

EEG data were recorded from 64 electrodes placed according to the extended 10-20 system (ActiveTwo, BioSemi, Amsterdam, The Netherlands). Four additional electrodes recorded vertical (vEOG) and horizontal (hEOG) eye movements. Preprocessing steps included high-pass filtering at 0.01 Hz (data of one participant in Experiment 1 were high-pass filtered at 0.5 Hz to remove excessive sweat artefacts), down-sampling from 2048 Hz to 512 Hz, and re-referencing to the common average. Eye blinks were removed from the data using adaptive spatial filtering based on individual blink templates computed from the vEOG (Ille, Berg, & Scherg, 2002). For the ERP analyses, the continuous data were cut into epochs from −50 to 600 ms relative to stimulus onset. All epochs were visually inspected for artefacts and artefactual trials were removed (on average 6.48% in Experiment 1 and 5.20% in Experiment 2). The artefact-free, epoched data were then low-pass filtered at 40 Hz and baseline corrected using a baseline from −50 to −5 ms. All data preprocessing and analyses were performed using SPM12 (www.fil.ion.ucl.ac.uk/spm, RRID:SCR_007037) and custom Matlab scripts (available on GitHub: https://github.com/PiaSchroeder/SomatosensoryTargetDetection_EEG). EEGLAB’s topoplot function was used to plot topographies (Delorme & Makeig, 2004, RRID:SCR_007292).

### Data analysis

#### Behaviour

Logistic functions were fitted to the behavioural data of each experimental block to obtain continuous models of the underlying psychometric functions (Wichmann & Hill, 2001). For each participant, the estimated slope and threshold parameters were then averaged across blocks to obtain one psychometric function per participant. An estimated detection probability <10% for intensity level 1 and >90% for intensity level 10 were defined as inclusion criteria. The rationale behind these criteria was that minimum and maximum detection probabilities outside these margins would indicate an incomplete sampling of the individual psychometric function (possibly due to changes in detection thresholds, response criteria, or erroneous reports). In Experiment 1 (direct report task), data of one participant and in Experiment 2 (matching task), data of three participants were excluded based on these criteria (**Figure 2**). To test whether the matching task successfully dissociated target detection from overt reports, Bayesian tests of association (Johnson & Albert, 1999) were performed on these variables for each participant in Experiment 2 and Bayes factors for independence are reported (BF01, i.e. Bayes factors in favour of the null hypothesis). Differences in reaction times between hits and misses were assessed using a Bayesian equivalent of the paired-sample t-test (Krekelberg, 2019) and Bayes factors in favour of a difference are reported (BF10). Following the recommendations by Kass and Raftery (1995), we consider 1 ≤ BF < 3 negligible, 3 ≤ BF < 20 positive, 20 ≤ BF < 150 strong, and 150 ≤ BF very strong evidence. All descriptive statistics are reported as mean ± standard deviation, except where otherwise noted.

#### EEG

EEG data of both experiments were analysed using Bayesian GLMs in combination with BMS. Seven GLMs were constructed that each contained an intercept regressor and a trial-wise experimental regressor: 1. intensity: the stimulus intensity levels presented on each trial (linear regressor), 2. P(detection): the trial-wise detection probability inferred from individual, block-wise psychometric functions (sigmoidal regressor), 3. detection: target detection, which was explicitly reported in Experiment 1 and inferred from match/mismatch reports and matching cues in Experiment 2 (binary regressor), 4. uncertainty: the slope of individual, block-wise psychometric functions modelling the expected uncertainty associated with target detection (inverse u-shaped regressor), 5. match: matches and mismatches between target detection and visual matching cues that were inferred from hit/miss reports and matching cues in Experiment 1 and explicitly reported in Experiment 2 (binary regressor), 6. cue: the white or dark grey matching cue, serving as a visual control model (binary regressor), and 7. null: a control model that contained the intercept regressor only to identify time segments that showed no effects. The seven GLMs were fitted to participant’s trial-wise EEG data using the Bayesian estimation scheme as implemented in SPM’s spm_vb_glmar.m function. This function approximates the posterior distributions of regression coefficients using variational Bayes (Penny, Kiebel, & Friston, 2003) and provides free energy approximations to the log model evidence (LME), which can be used for model comparison (Penny, Flandin, & Trujillo-Barreto, 2007). To obtain time-resolved estimates of LMEs for each model, electrode, and participant we fit our GLMs to each time point of each electrode individually. Thus, we set the autoregressive model order to zero, effectively reducing the error term to independent and identically distributed (IID) Gaussian errors. For the regression coefficients, we did not impose any constraints on the spatial continuity of effects and instead used a Gaussian prior with mean w0 = 0 and variance α^-1^ = 0.005 for all electrodes. All EEG data were z-scored across trials before model fitting to obtain data of comparable signal amplitudes.

The estimated LMEs were then used to perform BMS (BMS, Stephan et al., 2009) and obtain time-resolved exceedance probability (EP) estimates. Importantly, BMS allows comparing models that share variance to different degrees. If models are correlated, their shared variance reduces the relative difference between their LMEs. Accordingly, if correlated and uncorrelated models were included in the comparison individually, the correlated models would be at a disadvantage because they would have to compete against very similar models. At the same time, by assigning equal prior probability to correlated and uncorrelated models, identical portions of variance would be assigned too much prior weight, again resulting in an unfair comparison. BMS offers a simple solution to this problem by combining correlated models into a model family and adjusting the models’ prior probabilities accordingly (Penny et al., 2010). In our analysis, the intensity, detection probability, and detection regressors correlated positively, whereas all other regressors were expected to be uncorrelated. We thus combined the intensity, detection probability, and detection models into a model family (+family) and performed BMS on the family level. For time points that were best explained by the +family we further determined which of the individual +family models best explained the data by running the model comparison on the +family models only and weighting the resulting EPs by the +family EP.

Time points that yielded EPs ≥ .99 for any of the model families (*+family*, uncertainty, report, cue, null) were considered time points of interest. In addition, because BMS does not take into account the directionality of effects across participants (LMEs can be high regardless of whether an effect is positive or negative), we extracted beta estimates of the winning models’ experimental regressors and tested these for systematic deviation from zero using a Bayesian equivalent of the one-sample t-test (Krekelberg, 2019). This test ensured that only signals that systematically varied with any of our experimental regressors would be identified.

To further assess the influence of the different task requirements on detection-related ERPs, we extracted subsamples of hit and miss trials that were matched for stimulus intensity. For each participant, we identified all intensity levels that resulted in both hits and misses and randomly sampled trials such that the number of hits and misses was identical within each intensity level (**Supplementary Figure 1)**. The subsampled trials were then pooled across intensity levels (direct report task: 211.23 ± 47.35 trials per condition; matching task: 225.96 ± 43.32 trials per condition [mean ± standard deviation]) and grand-averaged hit and miss ERPs were plotted for electrodes of interest.

## Supplementary Figures

**Supplementary Figure 1.**
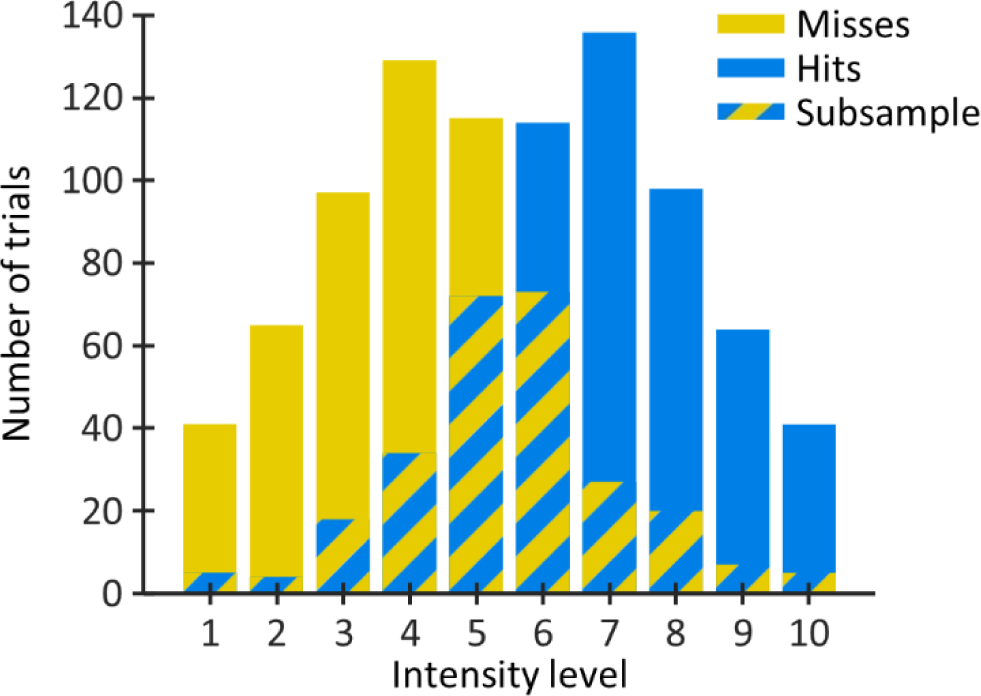
Sampling intensity-matched hit and miss trials. Trial distributions are shown for one exemplary participant. Lower intensity levels resulted in more miss trials (yellow) whereas higher intensity levels resulted in more hit trials (blue). To obtain intensity-matched subsamples of hit and miss trials, for each intensity level, we determined the number of trials obtained per condition and sampled as many trials from the condition with more trials as available for the condition with fewer trials. The subsampled trials (overlap) were then pooled across intensity levels to obtain a hit and a miss pool with identical intensity distributions.

**Supplementary Figure 2.**
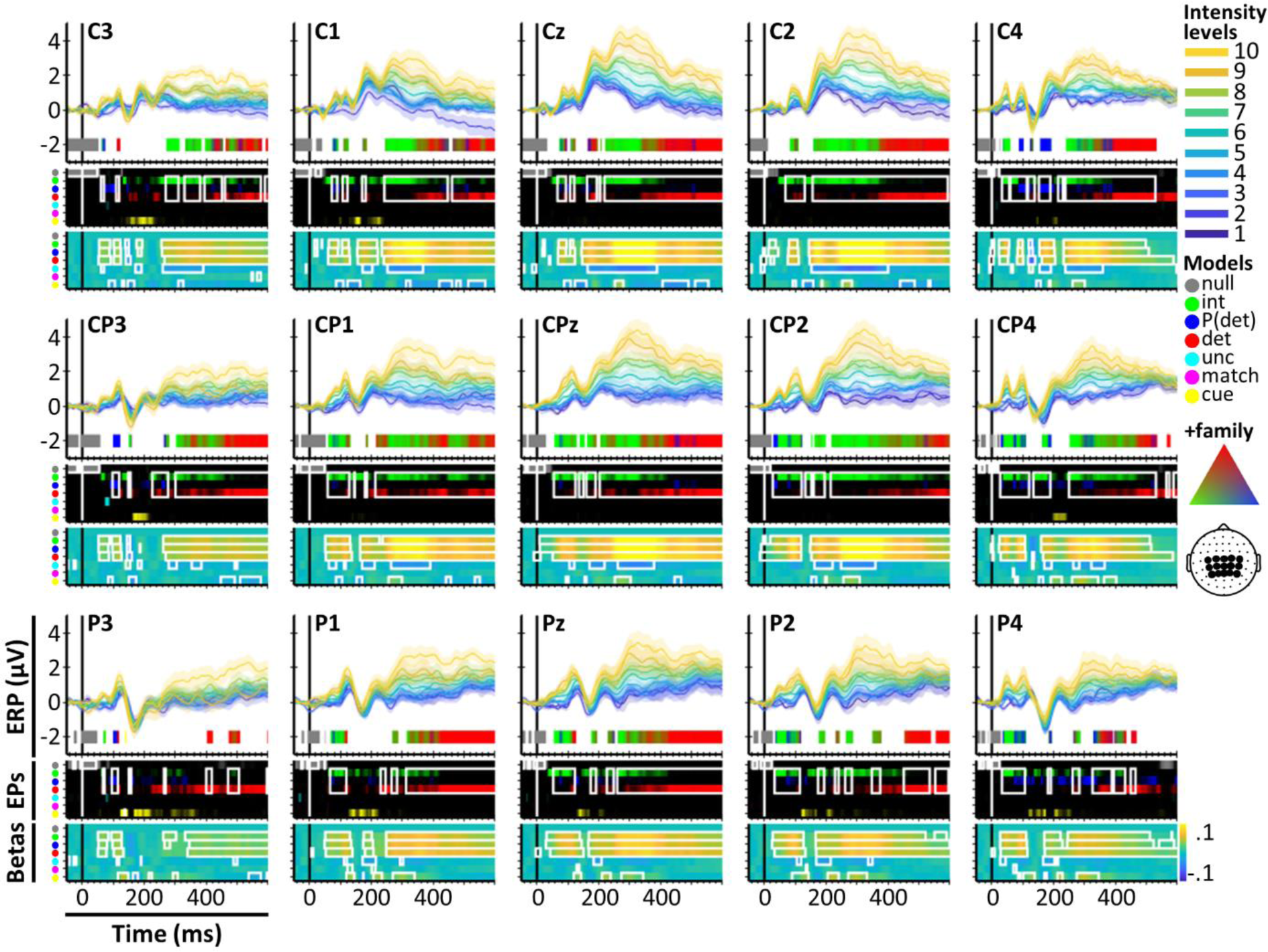
ERP and BMS results in centroparietal electrodes in the direct report task. Most electrodes show an early effect of stimulus intensity that transitions to an effect of target detection at ∼350 ms. Results are displayed as in Figure 3A. The scalp positions of the displayed electrodes are indicated in the schematic head on the right.

**Supplementary Figure 3.**
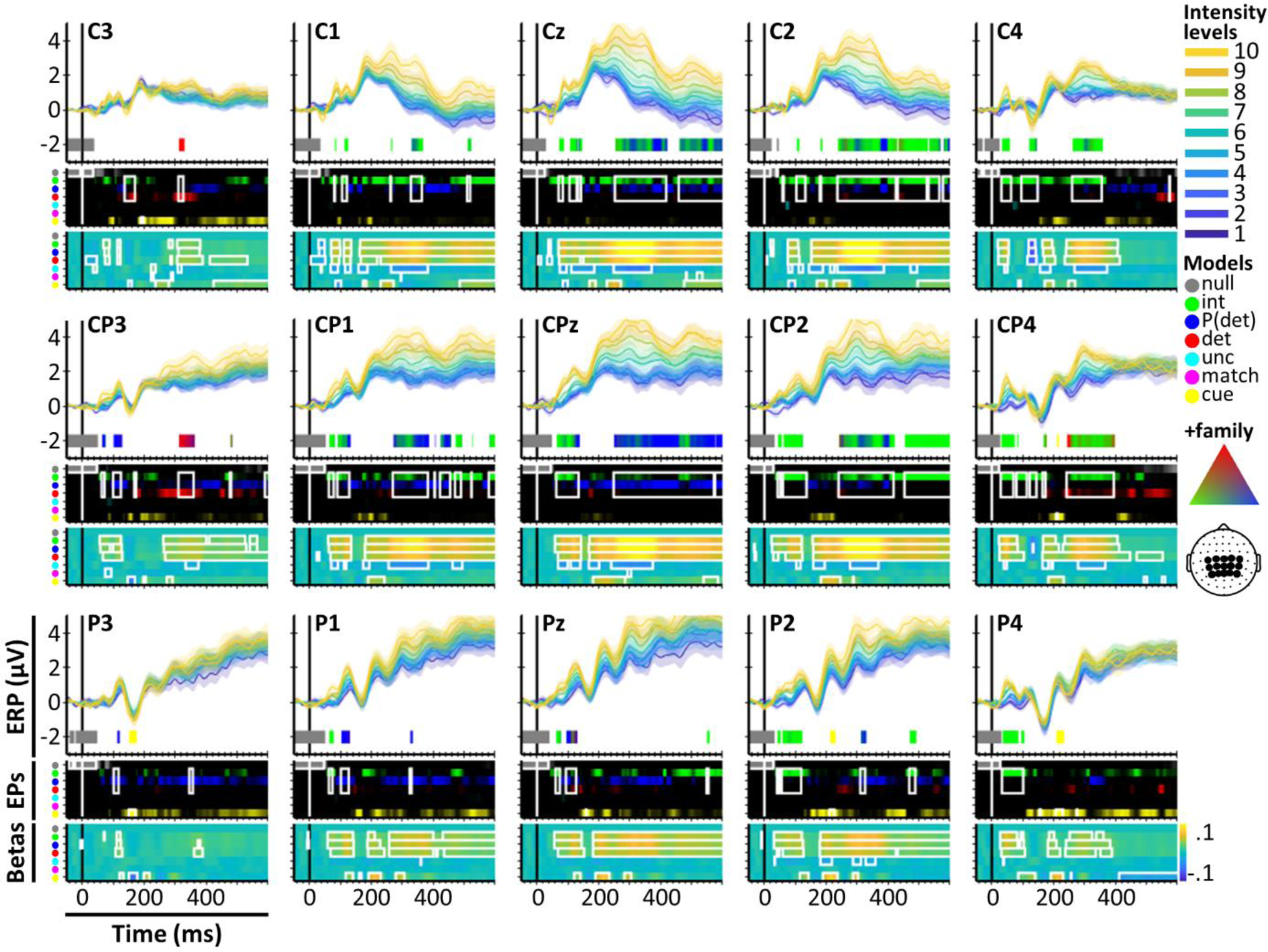
ERP and BMS results in centroparietal electrodes in the matching task. Most electrodes show effects of stimulus intensity or detection probability. The late detection effect observed in the direct report task is largely absent, with the exception of electrodes C3 and CP3, which show brief detection effects at ∼320 ms. Results are displayed as in Figure 3A. The scalp positions of the displayed electrodes are indicated in the schematic head on the right.

**Supplementary Figure 4.**
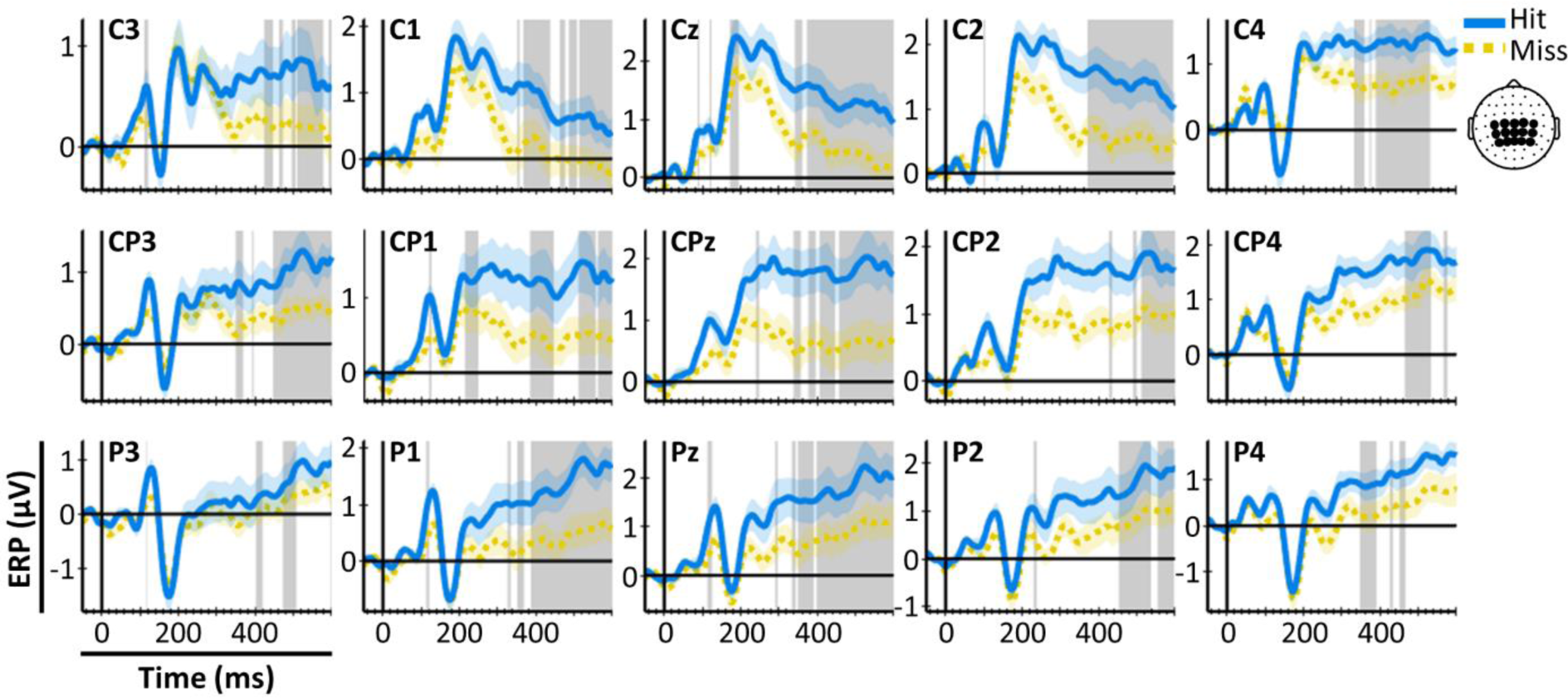
Intensity-matched hit and miss ERPs in centroparietal electrodes in the direct report task. Electrodes that were identified to show detection effects show a clear divergence of grandaveraged hit and miss ERPs in the P300 time window. Results are displayed as in Figure 5. The scalp positions of the displayed electrodes are indicated in the schematic head on the right.

**Supplementary Figure 5.**
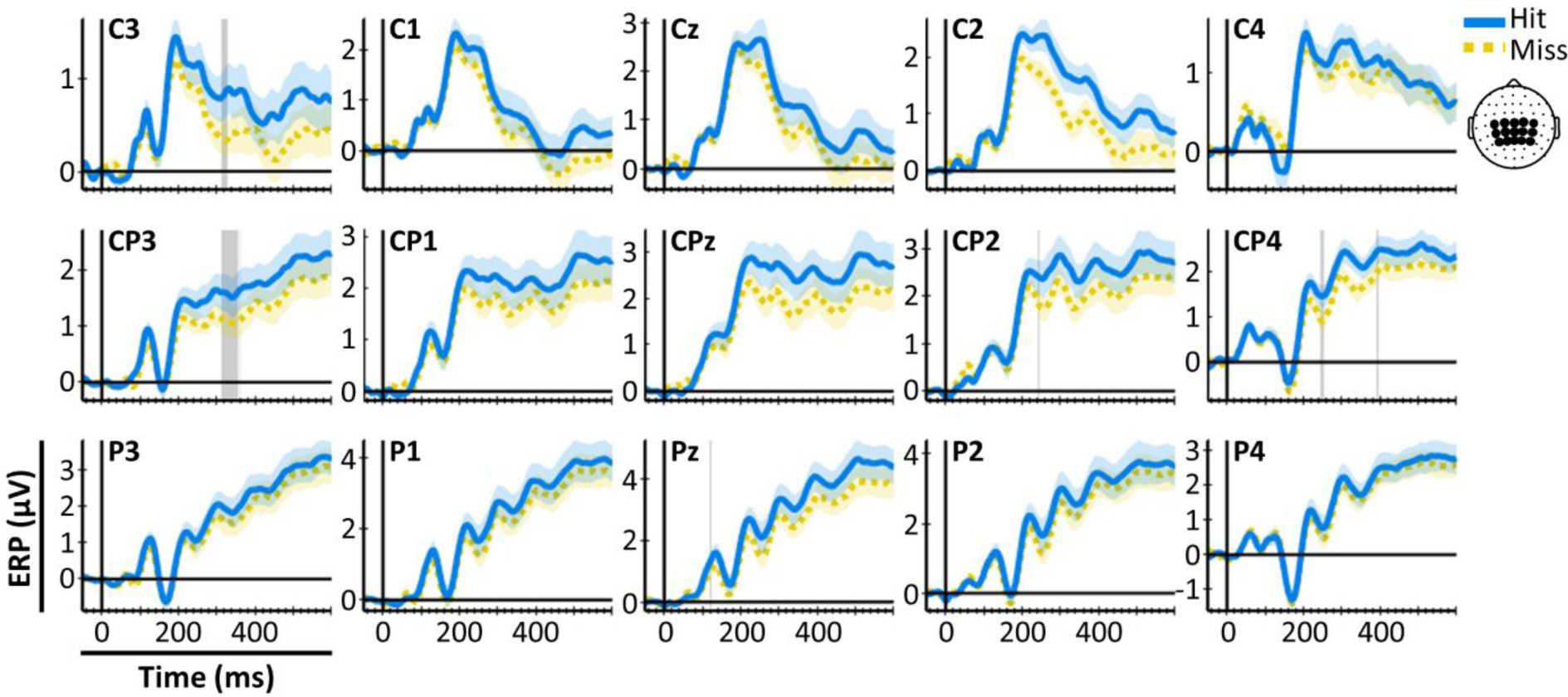
Intensity-matched hit and miss ERPs in centroparietal electrodes in the matching task. Grandaveraged ERPs for hits and misses are similar, even in electrodes that showed brief detection effects (compare electrodes C3 and CP3). Results are displayed as in Figure 5. The scalp positions of the displayed electrodes are indicated in the schematic head on the right.

